# Effect of natural precursors and micro/macroparticles addition on the morphology modulation of *Streptomyces toxytricini* KD18 stimulates lipstatin productivity

**DOI:** 10.1101/2023.07.02.547449

**Authors:** Khushboo, Namrata Dhaka, Kashyap Kumar Dubey

## Abstract

The cellular architecture of filamentous microbes is of great interest because it is frequently associated with secondary metabolite productivity and can be altered by cultivation conditions. Hence, the evaluation of cell morphology is of the utmost significance for better understanding of industrial processes involving filamentous bacteria. In the present study, effect of glass beads and silica particle addition have been seen in the enhancement of lipstatin production along with alterations in the morphology. The addition of glass beads and silica particles directed the morphology of *Streptomyces toxytricini* KD18 towards the formation of small pellets (0.3 to 0.4mm) with dispersed mycelia as compared to the control conditions (0.04 to 2mm). A four-fold increase in lipstatin production was achieved due to mechanical stress caused by glass beads and silica particles. The addition of natural precursors, i.e., *Aloe vera* pulp, *Hibiscus cannabinus* leaves and flowers improved the production of lipstatin from 2.06 mg/ml to 6.76 mg/ml, 11.4 mg/ml and 14.09 mg/ml, respectively along with alteration in the pellet morphology in 500 ml shake flask.

## Introduction

The complex architecture of actinomycetes is one of the most critical aspects to study. The strong correlation between the cellular morphology and yield of secondary metabolites is well known.^1–5^ The microscopic morphology of filamentous microbes is mainly the organization of small cylindrical fibers that divide to form branches and are termed hyphae. Under submerged conditions, the cellular architecture mainly depends on the culture conditions and ranges from loosely spread mycelium to a compact structure termed as pellet.^6^ The interaction between mycelia and the fragmentation process are the primary forces for driving macro-morphology evolution.^7^ Dense hyphal networks are known to affect the viability and productivity of mycelial pellets by restricting the passage of substrates to the center of the pellet.^8–10^ The cultivation of these organisms is difficult due to the variety of cell architecture. In recent years, different approaches came into picture for the tailoring of complex morphology to improve the production of secondary metabolites. These methods are collectively referred to as “morphology engineering”. It is well known that mechanical stress has a significant impact on the shape of filamentous cells.^11^

Apart from conventional approaches such as pH, temperature, agitation speed, medium composition, inoculum type and concentration, the most proven approach to alter the morphology of pellet is to change the osmolality of medium via addition of inorganic salts or micro/macroparticles for mechanical stress. These methods have already illustrated their utility and are currently awaiting more proof-of-concept studies. There is a strong link between cell shape and productivity in many filamentous bacteria.^12^ Depending on the strain and the product, microorganisms and their products can take on various morphological forms (mycelium, clumps, pellets). So far, there is no correlation between strain, genus, family, or product type that could help us to better understand how filamentous production systems work.^1, 13^

The aim of the present study was the morphological investigation and improvement of lipstatin production via increasing mass transfer in the pellet of *S. toxytricini* KD18 cultivated under different culture conditions. Effect of leucine rich natural precursors such as *Aloe vera* pulp, *Hibiscus* leaves and flowers was examined in the present study. The combined effect of these natural precursors on the morphology and secondary metabolite production was also examined at both, the shake flask and lab-scale bioreactor level.

## Materials and Methods

### Microbial strain and culture conditions

*S. toxytricini* KD18, previously isolated from industrial soil ^14^ was used in the present study. The strain was cultivated on YMG (yeast extract, malt extract and glucose) agar plates and incubated at 28±2 °C for 7 days. The morphology of the pellet was examined in the growth medium containing yeast extract (4 g/L), malt extract (10 g/L), glucose (4 g/L) and CaCO_3_ (2 g/L). Single colony from agar plates was inoculated in the growth medium and incubated at 28°C and 200 rpm for 24 h to form the primary culture. The primary culture was inoculated into the fresh growth medium to form the secondary culture and incubated under the same culture conditions to analyse the pellet architecture.

### Optimization of growth parameters for pellet morphology

The effect of physical culture parameters on the growth and morphology of *S. toxytricini* KD18 was examined at variable pH (5-8) and temperature (25°C, 28°C, 30°C and 35°C). Similarly, influence of (i) different agitation speed (100-300 rpm) and (ii) percent of inoculum (1-6%) were also examined. Various carbon sources (sucrose, maltose, fructose, lactose, D-mannitol and starch instead of dextrose) and nitrogen sources (peptone, soya peptone, sodium nitrate, casein and ammonium nitrate instead of yeast extract) at 4 g/L concentration were screened to optimize the growth parameters. Different inorganic salts i.e., MgSO_4_.7H_2_O, CuSO_4_, ZnSO_4_.7H_2_O and FeSO_4_.7H_2_O were utilized to grow the bacterial culture by replacing CaCO_3_.

### Micro/macroparticles cultivation

The optimized parameters providing maximum biomass were applied to examine the effect of micro/macroparticles in pellet formation. The impact of silica oxide (SiO_2_) particles and glass beads on pellet morphology and lipstatin production was examined. The culture was inoculated with 2% of inoculum. In order to carry out particle-based cultivations, 5 g/L fractured, porous L60 SiO_2_ particles (40-60 µm size) were used. To examine the impact of glass beads, 2% inoculum was seeded in 250 ml flask with 100 ml YMG medium and 5 g glass beads (2.5-5 mm diameter) followed by incubation at 28°C with 200 rpm.

### Impact of natural precursors on lipstatin productivity

Naturally available, *Hibiscus cannabinus* leaves (HLs) (1.5 g/100 ml), flowers (HFs) (1.5 g/100 ml) and *Aloe vera* pulp (5 ml/100 ml) were also used as medium components to study their effect on the pellet morphology and lipstatin production.

### Culture conditions at shake flask level

Culture conditions were adopted from Kumar and Dubey, 2017.^15^ One flask had the production medium (PM) was enriched with *Aloe vera* pulp while another flask had PM supplemented with HLs and third flask had HFs in the PM. The PM was also enriched with combined supplementation of *Aloe vera* pulp, HLs and HFs. Culture was examined for lipstatin production at 264 h of incubation by HPLC analysis.

### Fermentation conditions at 5 L bioreactor level

Experiments were carried out in 5 L fermenter with 3 L working volume. The culture medium for lab-scale bioreactor was commercial type containing glycerol 22.5 ml/L, soy oil 25.0 ml/L, soy flour 35.0 g/L, soy lecithin 15.0 g/L, polypropylene glycol 0.5 g/L, leucine 50 mM, yeast extract 4.0 g/L, calcium carbonate 2.0 g/L. The fermentation broth was inoculated with 10% seed medium from shake flask. The pH of culture medium was maintained using 1N NaOH and 1N HCl solutions and incubation conditions were same as that of shake flask. Similar to the shake flask, the PM was also enriched with combined supplementation of pulp, HLs and HFs at the bioreactor level.

### Extraction of lipstatin

The extraction method was proposed by Kumar et al., 2012 in which extraction of fermentation broth was performed three times with acetone: hexane (3:2). The resulted organic layers were combined and passed through sodium sulphate placed over cotton using conical funnel. The resulting organic layer was concentrated to form an oily liquid which was further dissolved in hexane. This crude lipstatin was subjected for HPLC analysis.^16, 17^

### Morphological analysis

For morphological analysis, methylene blue staining was performed. The methylene blue stain was poured on the heat fixed smear of *S. toxytricini* KD18 strain on the glass slides and left for 5 min to air dry.

## Results

### Morphological analysis of pellet

Morphological examination of *S. toxytricini* KD18 strain was performed via methylene blue staining and visualization was done under inverted microscope. Formation of interconnected network of mycelia i.e., pellet in the submerged culture is one of the main characteristics of *Streptomyces* (Fig. 1).

**Fig. 1.**
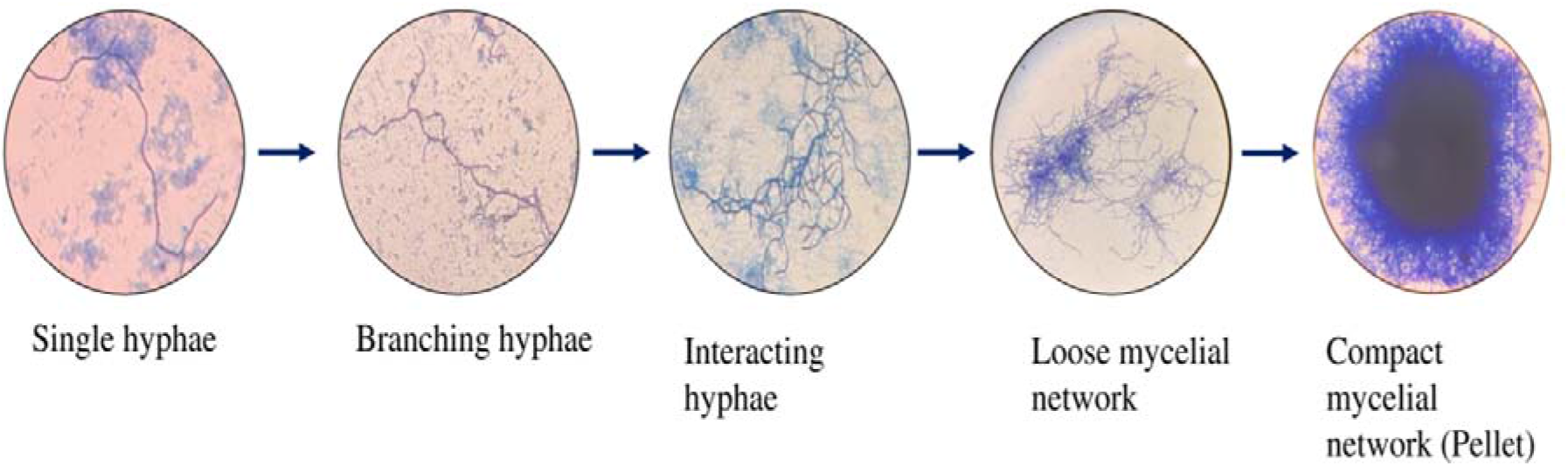
Steps involved in pellet formation. Diffused mycelia grow through the hyphal tips, which form the mycelial branches. The mycelial branches get interconnected with each other and form clumps. The mycelial clumps become more condensed and form the pellet.

### Optimization of inoculum size

Here, 1-6% (v/v) inoculum size was used to examine the effect on the biomass formation. The biomass formation enhances with enhancement in inoculum size and maximum biomass (7.9 mg/ml) was formed at 6% inoculum with 35 µm-2 mm pellet size having well developed mycelia at the periphery (Table 1). It was observed that the pellet diameter and inoculum size are inversely proportional to each other i.e., increase in the inoculum volume resulted in smaller pellet formation. The biomass produced at 5% inoculum was nearly similar to 6% inoculum size and the pellet diameter was up to 1mm which might be a better condition for improved production of secondary metabolite.

**Table 1.**
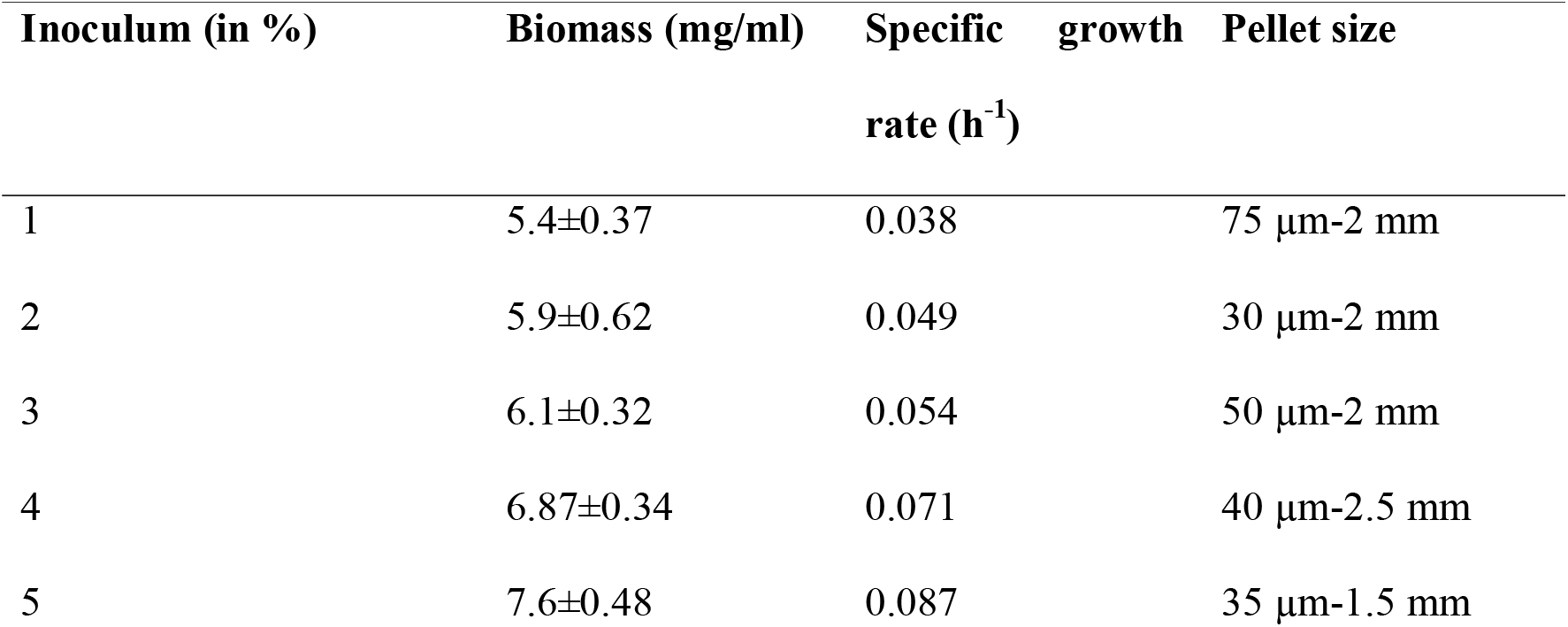
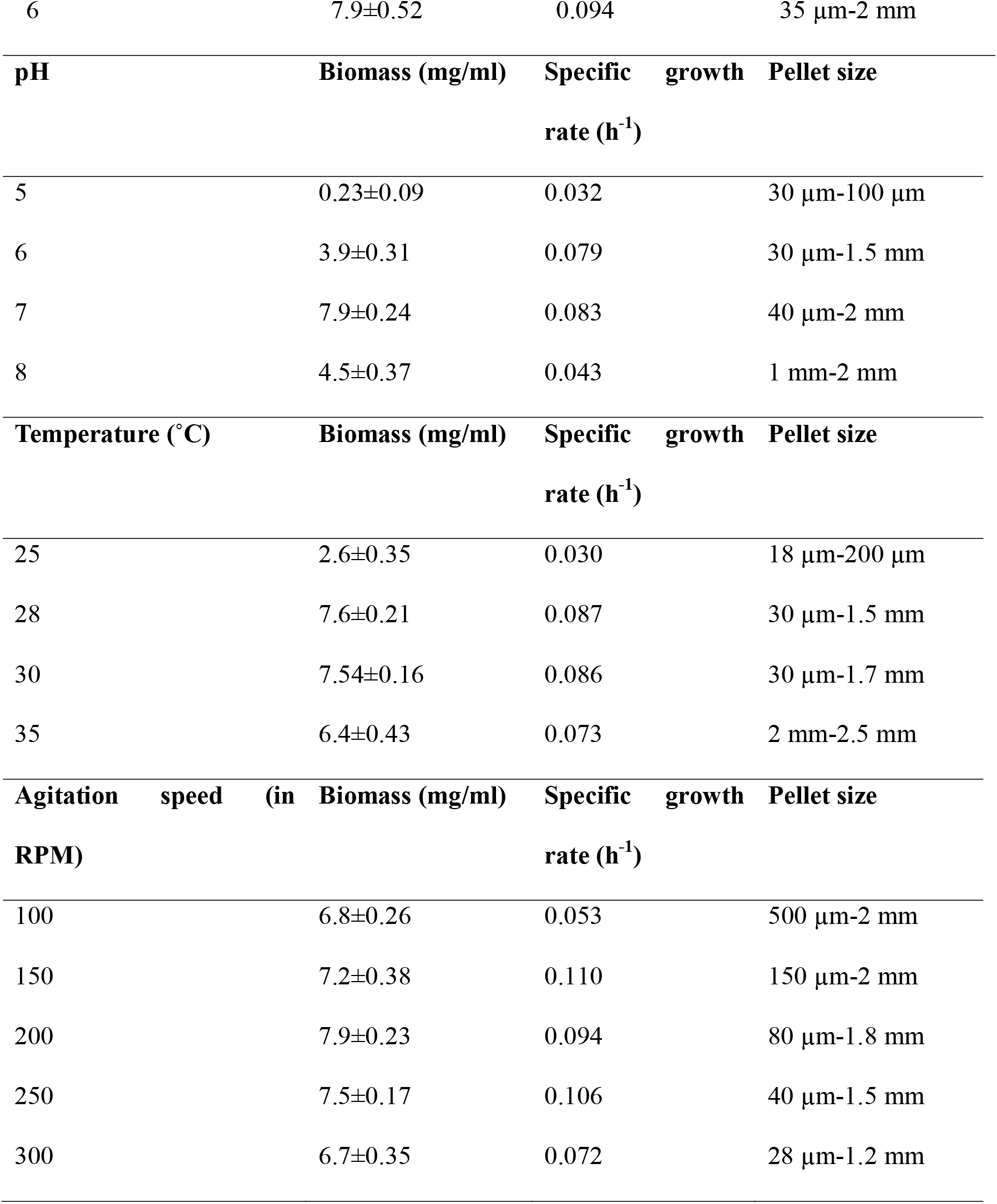
Impact of different physicochemical parameters on biomass and pellet formation. Control conditions: Inoculum-2%; pH-7; Temperature-28 °C; Agitation speed-200 rpm.

### Optimization of pH

It is well known that the enzymes work at optimum pH conditions. Similar observation was made in the case of *Streptomyces*. In the present study, effect of different pH conditions (5-8) was examined over the growth and morphology of *S. toxytricini* KD18. It was observed that the biomass as well as pellet diameter of the culture increased along with increase in the pH. At pH 5 very small size pellets (30µm-100µm) were observed. The small size pellets are better for the production of secondary metabolites but along with pellet size, biomass is also an important factor. For small pellet formation pH 5 was good but the biomass formation (0.23 mg/ml) was also very less. 7.9 mg/ml was the maximum biomass produced at pH 7.5 with 35 µm-2 mm of the pellet (Table 1).

### Optimization of temperature

In the current investigation, the growth of isolated strain KD18 was examined at 25, 28, 30 and 35°C temperature conditions. Dispersed mycelial growth was observed at 28°C with 7.6 mg/ml biomass and 30 µm-2 mm size of pellet formation. Scanty and late growth was observed at 25°C with small size of pellet formation. The diameter of pellet increased with increase in temperature. Very large size of pellets (2 mm-2.5 mm) were formed at 35°C without any dispersed mycelia. Growth and morphology of pellet were nearly similar at 28 and 30°C (Table 1).

### Optimization of agitation speed

High agitation speed means high shear stress which leads to the mechanical fragmentation of hyphae. As a result, pellet with small diameter would be formed.^18–20^ Similar effect of agitation speed was observed in the present study. As the agitation speed increased, significant reduction in the diameter of pellet was observed. At 300 rpm, the size of pellet was in between 28 µm-1.2 mm while at 100 rpm, the pellet size was up to 2 mm. However, the biomass production was not remarkably affected by the agitation speed. Nearly, same amount of biomass (6.8-7.9 mg/ml) was produced under all agitation speeds applied in this study (Table 1).

### Optimization of carbon source

KD18 strain was grown in the medium with different forms of saccharides such as dextrose, fructose (monosaccharide), maltose, sucrose, lactose (disaccharide) and mannitol, starch (polysaccharide). It was observed that dextrose supported the maximum growth of KD18 with 53.6 mg/ml biomass formation and the specific content of lipstatin was 38.43 mg/g of DCW at 264 h of incubation (Fig. 2). Fine mycelial formation was observed under the effect of all carbon sources and nearly same size of pellet was formed without any discernible impact from the various carbon sources. Minimum biomass i.e., 23.2 mg/ml was formed in the growth medium supplemented with sucrose at 264 h of incubation where the diameter of mycelial clump was 35μm-1.5mm. In dextrose enriched medium, 30 μm-2 mm diameter of pellet was recorded. Overall, the pellet obtained with all the carbon sources ranged in size from 15 μm-2 mm with better mycelial growth at the periphery.

**Fig. 2.**
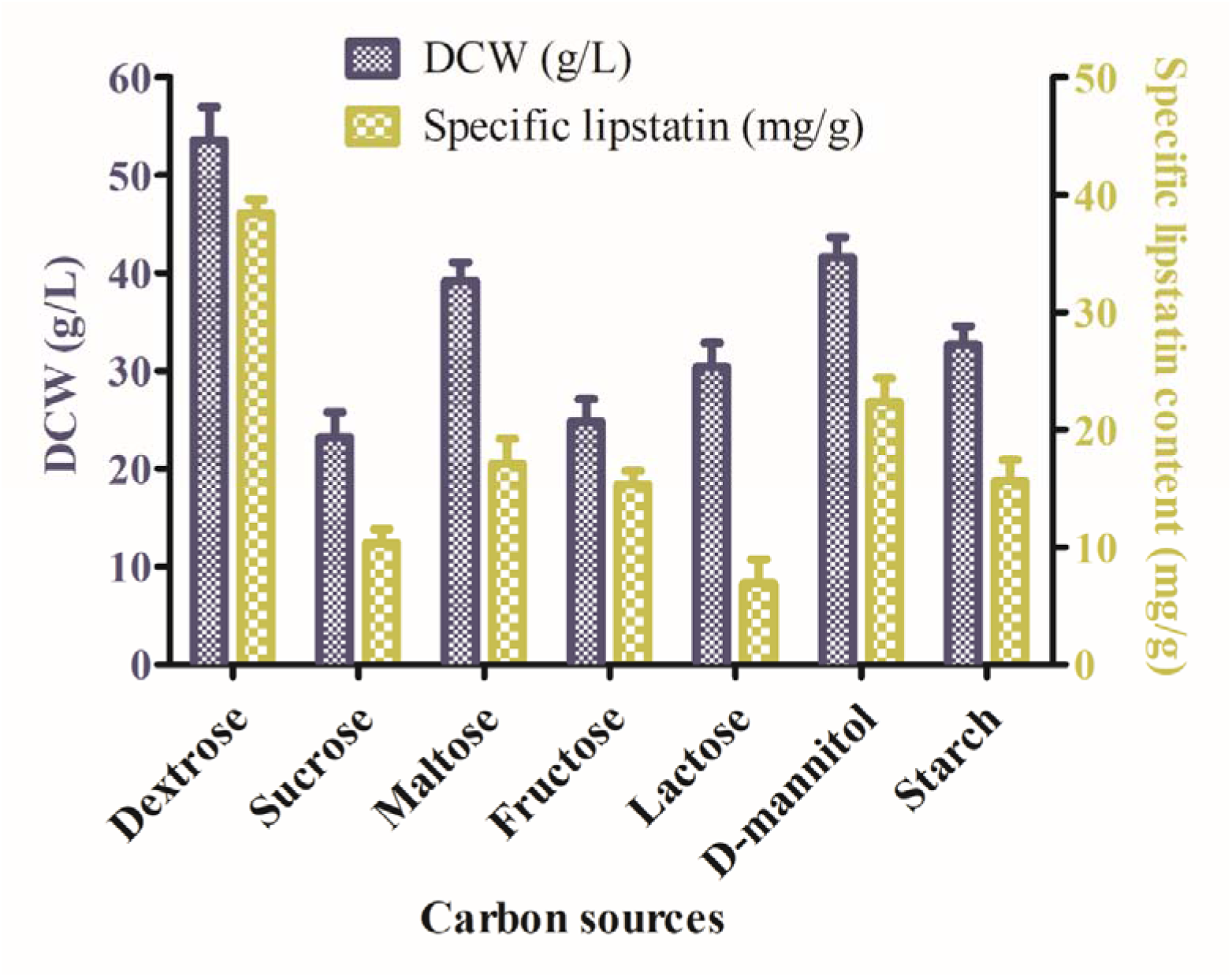
Growth of *Streptomyces toxytricini* KD18 with different carbon sources. Different amount of lipstatin was produced with different carbon sources. Maximum amount of lipstatin production was observed with dextrose (control) whereas sucrose was least favorable for lipstatin production as well as for the growth. Experiments were performed in triplicates and bar represents the mean ±SD.

### Optimization of nitrogen sources

In the present study, impact of various nitrogen sources on the pellet morphology, biomass formation and secondary metabolite production was examined. Yeast extract supplemented medium (control) produced higher biomass i.e., 63.2 mg/ml with 40μm-1.58mm pellet formation at 264 h of incubation. Lowest biomass (28 mg/ml) was produced in the medium supplemented with ammonium sulphate while smallest size of pellet was formed in the medium enriched with sodium nitrate (45 μm–350 μm) and ammonium sulphate (28 μm–400 μm). However, well developed mycelia were observed around the periphery of the pellet formed with (NH_4_)_2_SO_4_ and NaNO_3_ enriched medium. The specific lipstatin content at 264 h of incubation with different nitrogen source is represented in fig. 3.

**Fig. 3.**
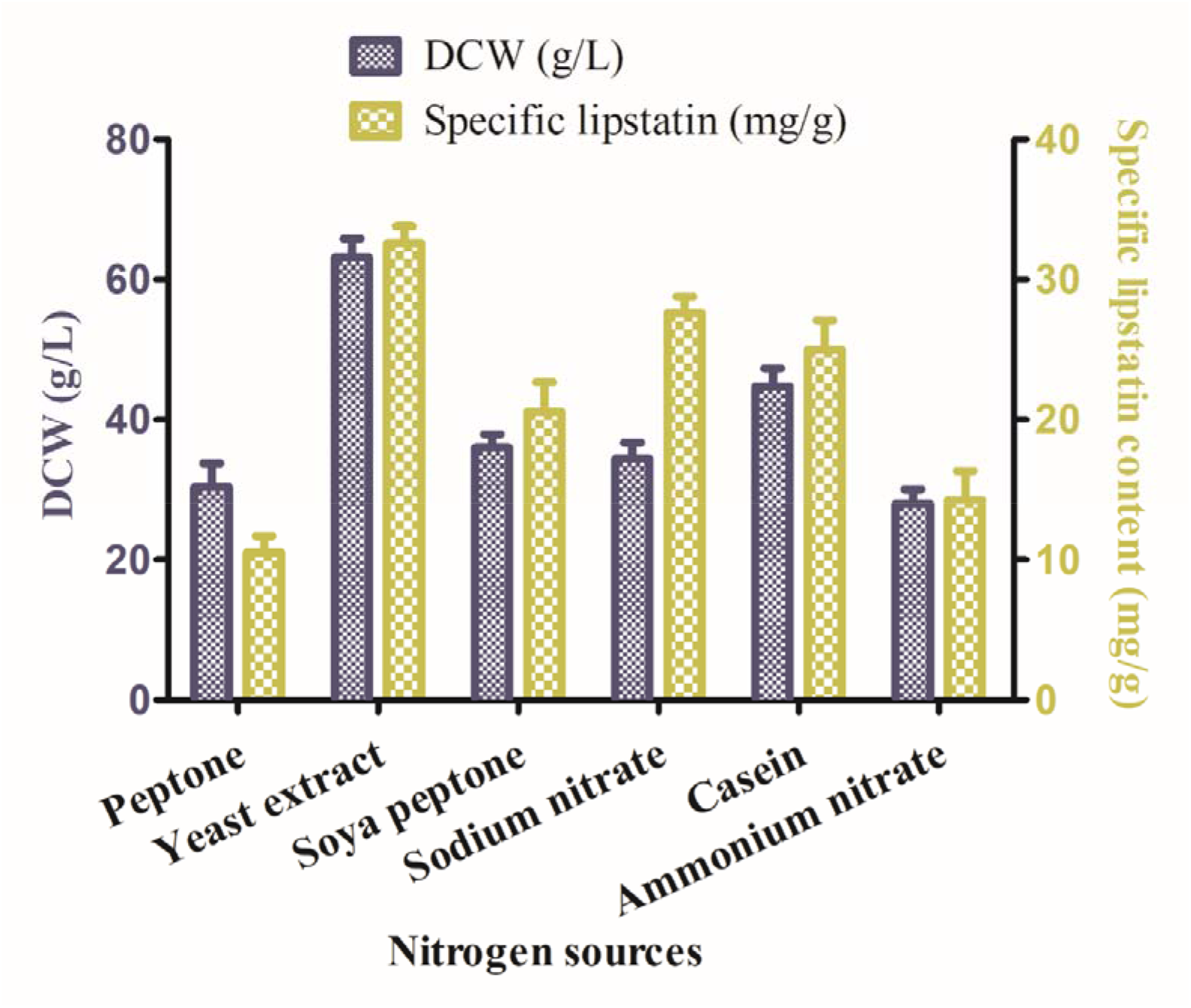
Growth of *S. toxytricini* KD18 with different nitrogen sources. Maximum lipstatin production was achieved with yeast extract (control). Ammonium nitrate and peptone were not effective for the lipstatin production.

### Optimization of inorganic salts

The divalent ions used in the present study showed significant effect on the biomass formation. The data presented in the fig. 4 revealed that *S. toxytricini* KD18 grew well in calcium and magnesium supplemented medium.

**Fig. 4.**
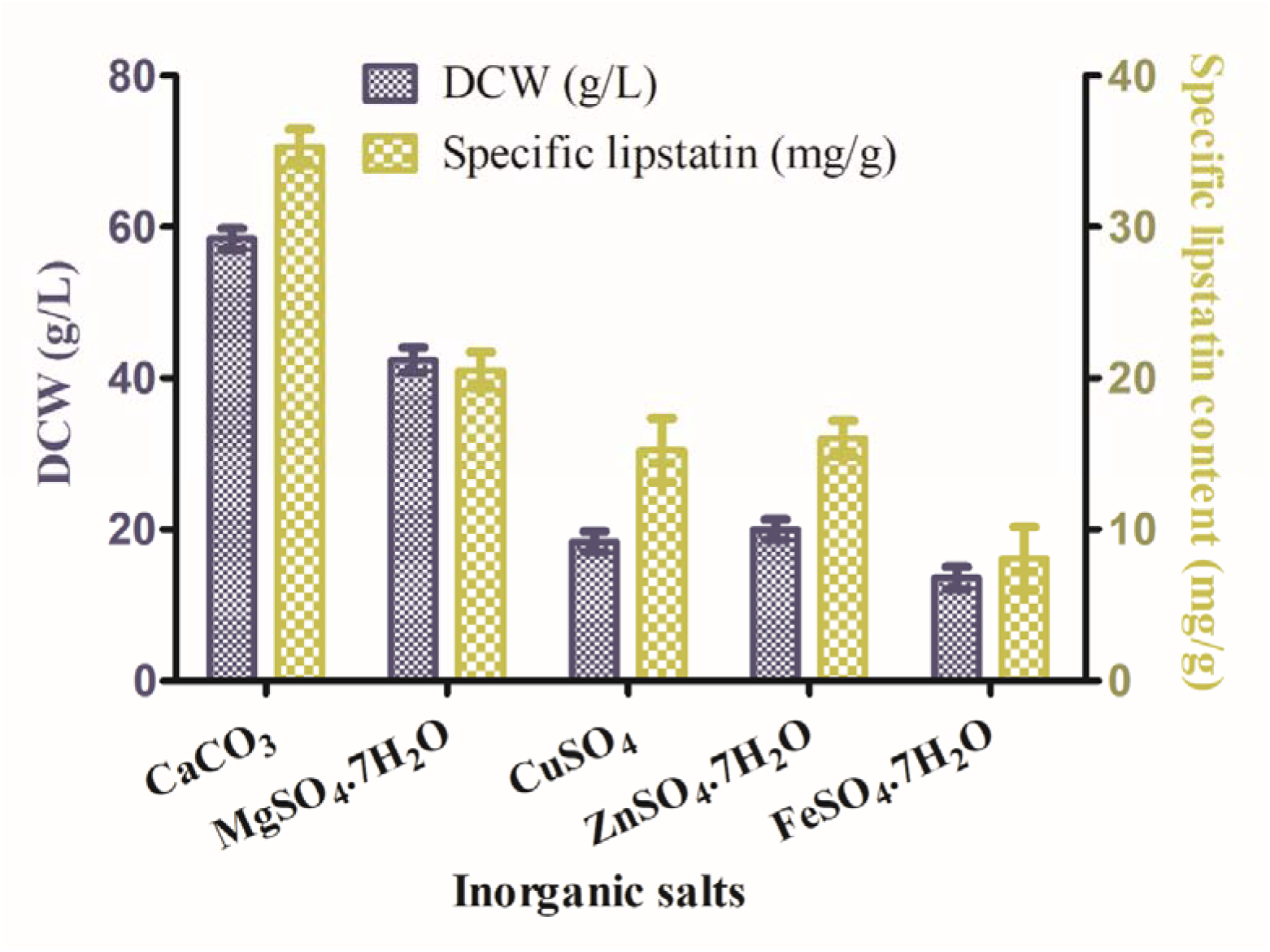
Effect of different inorganic salts on the growth and lipstatin production. CuSO_4_, FeSO_4_, ZnSO_4_ were showing adverse effect on lipstatin production as compared to control (CaCO_3_).

The maximum biomass i.e., 58.4 mg/ml was recorded in the medium containing CaCO_3_ with 35 µm-2 mm pellet formation. Zn^2+^, Cu^2+^ and Fe^2+^ showed negative effect on the cell growth as very less biomass was obtained in the medium supplemented with CuSO_4_, ZnSO_4_ and FeSO_4_.7H_2_O, respectively. The specific content of lipstatin was also very less with the addition of Zn^2+^, Cu^2+^ and Fe^2+^ salts in the PM (Fig. 4). The pellet architecture was also affected by these supplements as no mycelial growth was observed on the pellet periphery. Diffused mycelia were observed in growth medium enriched with FeSO_4_.7H_2_O.

From the above optimized parameters, further study was performed to examine the production of lipstatin. Here, dextrose and yeast extract were used as carbon and nitrogen source, respectively. The culture was grown in PM with 7.5 pH at 28°C and 200 rpm. These conditions were used as the highest biomass was formed along with normal morphology of pellet.

### Pellet morphology and lipstatin production

Under normal PM (control), 2.06 mg/ml lipstatin production was achieved whereas the diameter of pellet was 40µm-2mm (Fig. 5A).

**Fig. 5.**
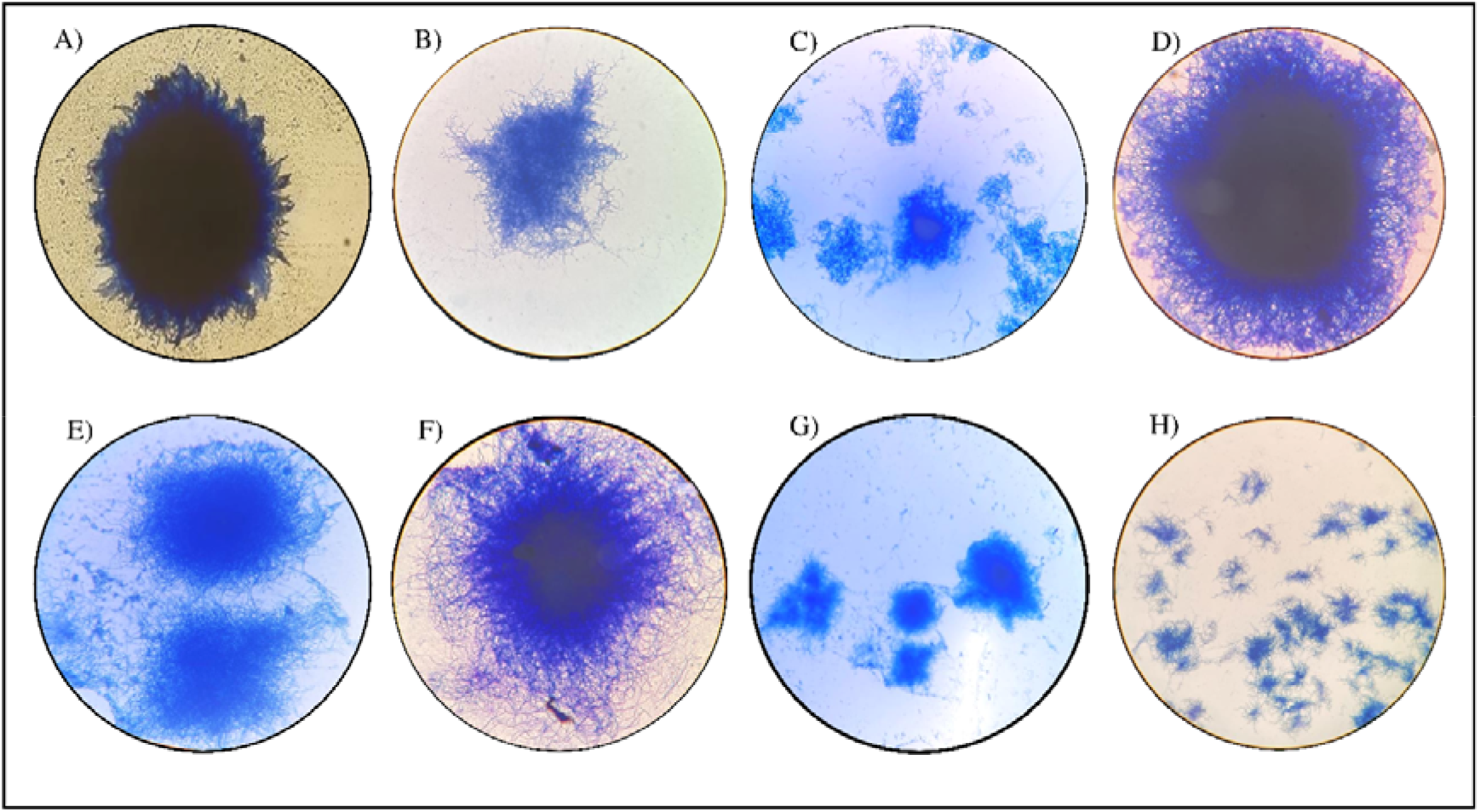
Representatives of pellet formation under different conditions. (A) PM (control); PM supplemented with (B) glass beads; (C) silica particles; (D) *Aloe vera* pulp; (E) HLs; (F) HFs; (G) Pulp+ HLs+ HFs (Flask); (H) Pulp+ HLs+ HFs (Bioreactor). Scale bar-200µm for each figure.

### Effect of glass beads

The mechanical shear force to the pellet of actinomycetes might be impaired through glass beads at the shake flask cultures. Here, the collisions among the medium particles, glass beads and mycelia or hyphae leads to the fragmentation which in turn alters the morphology. In the present study, addition of glass beads to the PM significantly enhanced the production of secondary metabolite. The shear stress caused by the glass beads resulted in the formation of very small size pellet (300 µm) with well-developed mycelia at the periphery (Fig. 5B). In addition, dispersed mycelia were also formed. The production of lipstatin was improved upto 4 folds as compared to the control while the diameter of pellet was reduced upto 15%.

### Effect of silica particles

In the present study, addition of silica particles remarkably affected the morphology as well as lipstatin production. The pellets were formed in the range of 25-400 µm, significant reduction in the size as compared to control (40 µm-2 mm) (Fig. 5C). Similar to the addition of glass beads, dispersed mycelia were also observed in the culture. 4.5-fold enhancement in lipstatin production was achieved via the addition of silica particles.

### Effect of natural precursors

As mentioned in the literature, *Aloe vera* pulp, HLs and HFs are rich sources of leucine (an important component for lipstatin biosynthesis) isoleucine, phenylalanine and many other amino acids.^21, 22^ Supplementation of these components to the *Streptomyces* culture proved effective for the pellet morphology as well as lipstatin production. However, no other study was reported related to the effect of *Aloe vera* pulp, HLs and HFs on the pellet morphology and secondary metabolite production.

Medium enriched with *Aloe vera* pulp formed similar size of pellet to that of the control i.e., 50 µm-2 mm (Fig. 5D). However, the production of lipstatin was higher i.e., 6.76 mg/ml in pulp supplemented medium which was 3 folds higher as compared to control. The results were in contrast with the results of silica particles and glass beads. In case of micro and macroparticles, reduction in the pellet diameter with enhanced production of lipstatin was observed which indicates that the smaller pellet yields higher lipstatin.

The micro and macroparticles were not adding any nutrient value to the PM as a result only the morphological effect was involved in enhanced production of lipstatin. On the other hand, *Aloe vera* pulp was affecting the morphology of pellet along with improvement of nutritional value. So, here the factor of nutrients supply comes into limelight which put major effect on the secondary metabolite production. The supplementation of *H. cannabinus* leaves and flowers in the PM formed 30µm-1mm and 25 µm-1 mm size of pellet, respectively (Fig. 5E and 5F). 5.5- and 6.8-fold increment in the yield of lipstatin was achieved via the medium enriched with *H. cannabinus* leaves and flowers, respectively. Here, the size of pellet reduced in to addition to the improvement in the nutrient value of medium.

Enrichment of PM with pulp, *H. cannabinus* leaves and flowers was highly effective for the lipstatin production as 14.79 folds’ increment in lipstatin production was gained at the shake flask level. The diameter of the pellet was in the range of 45 µm-500 µm which was also significantly smaller as compared to the control (Fig. 5G).

### Lipstatin production at bioreactor level

In order to validate lipstatin production, *Streptomyces* culture was grown at the lab-scale bioreactor with the same culture conditions. Significant increment in the production of lipstatin was observed in 5 L bioreactor. Highly dispersed morphology was observed (Fig. 5H). 37.97 mg/ml of the lipstatin was produced at the fermenter level which was 18.4 folds higher as compared to the shake flask culture (Table 2).

**Table 2.** Effect of micro/macroparticles and other supplements on pellet formation and lipstatin production.

| **Types of medium** | **Pellet diameter** | **Lipstatin**<br>**Production (mg/ml)** |
| --- | --- | --- |
| PM (Control) | 40 μm-2 mm | 2.06 |
| PM+ glass beads | 300 μm with dispersed morphology | 8.12 |
| PM+ silica particles | 25 μm-400 μm along with dispersed morphology | 9.19 |
| PM+ *Aloe vera* pulp | 50 µm-2 mm | 6.76 |
| PM+ HLs | 30 µm-1 mm | 11.4 |
| PM+ HFs | 25 µm-1 mm | 14.09 |
| PM+ Pulp+ HLs+ HFs (Shake flask) | 45 µm-500 µm | 29.58 |
| PM+ Pulp+ HLs+ HFs (5 L Bioreactor) | 100 µm with dispersed mycelia | 37.97 |

## Discussion

The growth of microbes is highly dependent on the medium components. Different carbon, nitrogen, salt, and mineral sources have unlike impact on the microbial growth.^23–25^ The microbes are able to grow on any type of medium but in order to obtain higher biomass, most suitable growth medium needs to be provided.

Minimum biomass i.e., 2.9 mg/ml was formed in the growth medium supplemented with sucrose. Similar findings were reported by Jonsbu et al., 2002 where glucose favoured the growth of *S. noursei* while less biomass formation was observed with sucrose and fructose supplemented medium.^26^ Different *Streptomyces* sp. prefer different carbon sources for their growth and secondary metabolite formation. In various *Streptomyces*, glucose favors the growth while represses the secondary metabolite production as observed in case of *S. antibioticus* where addition of glucose adversely affects the production of actinomycin.^27^ However, for most of actinomycetes, glucose works in a positive manner for the growth. *S. clavuligerus* is an exceptional case which utilizes glycerol instead of glucose for growth.^28^

Catabolic inhibition of secondary metabolite might be linked with the inactivation of necessary enzymes linked with secondary metabolite formation. In case of *Nocardia lactamdurans*, glucose-6-phosphate inactivates deacetoxycephalosporin C synthase.^29^ *S. halstedii* utilizes maltose as a sole carbon source for the normal growth. In another study it was observed that monosaccharides were the most favored carbon source utilized by the isolated strains of actinomycetes from water sources.^30^

In case of inorganic salts, Leung et al., 2006 observed similar result in context of MgSO_4_ supplemented medium for the rapid and normal growth of *Tolypocladium* fungus ^31^ Along with change in the osmolarity, minerals are essential for the growth and multiplication of microbes, and they also act as cofactors for regulation of various catalytic activities. However, higher concentration of these inorganic salts might be toxic to the cell and they should therefore be used in small amount.

As the integral part of DNA as well as proteins, nitrogen is principal element for all living systems.^32^ Metabolism of nitrogen along with its scavenging process is very much essential for saprophytic microbes like *Streptomyces*. In addition, nitrogen is the part of various secondary metabolites which might be important for the adaptation of microbes to their surroundings.^33^ The result of the present study were in contrast to the study performed by Ababutain et al., 2013 on *Streptomyces* sp. MS-266 Dm4 where NaNO_3_ was suitable for high biomass formation.^34^ Jonsbu et al., (2000) also examined the effect of nitrogen sources on the growth and secondary metabolite formation and observed similar results as that of present study. Supplementation of NaNO_3_ increases the duration of lag phase as compared to other nitrogen sources resulting in less biomass formation along with reduced pellet diameter.^35^

As mentioned above, the temperature conditions are different for different *Streptomyces* species. Thakur et al. (2019) observed that the strain isolated from tea garden soil, *Streptomyces* sp. 201 preferred 35°C for better mycelial growth while the production of secondary metabolite was maximum at 30°C.^36^ In another study, *S. californicus* produced highest amount of griseorhodin at 28°C.^37^ Similar findings were reported by Kokare et al., 2007 where the isolated strain *Streptomyces* S1 produced secondary metabolite (bioemulsifier) at 28°C.^38^

In case of pH optimization, the findings were similar to that of Thakur et al., 2009 where *Streptomyces* sp. 201 was able to grow at pH 7-8. In another study, *S. thermoviolaceus* required pH 7 for normal growth.^39^ *S. coelicolor* also produced actinorhodin at pH 7.^36, 40^ Similar influence of pH was observed on the fungal morphology examined by Zhou et al., 2000, where diameter of pellet decreased with decrease in pH.^41^ It has been suggested that the negative or positive charge on the cell wall surface of microbes alters with alteration in the pH conditions which causes repulsion and restricts the attachment between mycelia.^42^ However, the optimum pH for the mycelia or pellet formation varies with different microbes depending on their surface charge.^43, 44^

Agitation might affect the microbes in a variety of ways including alteration in morphology, biomass formation and secondary metabolite production along with damage to the cell structure.^45, 46^ Zhou et al., (2018) described that the maximum biomass of *S. kanasenisi* ZX01 was formed at 200 rpm which was slightly higher than the biomass formed at other agitation conditions. However, increase in agitation speed up to 300 rpm imposed unbearable mechanical force to the cell and resulted in cell damage.^47^ In another study, high rotatory speed adversely affected the growth of *Streptomyces* sp. P6621-RS1726.^48^

The results obtained for inoculum size optimization were similar to the study described by Glazebrook and Vining, 1992, where *S. akiyoshiensis* formed small size of pellet when cultivated with inoculum of high percentage.^49^ Domingues et al.,1999 also observed similar results in case of *Trichoderma reesei* where pellet growth was higher in the medium inoculated with less inoculum percentage.^50^ Similar findings were described by Dobson et al., 2008 where 88% increment in geldanamycin production and 70% reduction in the pellet diameter of *S. hygroscopicus* was achieved.^19^

The results reported by Yepes-García et al., 2020 were contradictory to the observations of present study. However, only 0.4 mm size glass beads were used to examine the morphological differentiation of *S. clavuligerus*. Very small size of glass beads might be the reason for no effect on morphology as well as production of Clavulanic Acid.^51^ Sohoni et al., 2012 examined the effect of glass beads on the morphology of *S. coelicolor* grown in microtiter plate and witnessed the dispersed and reproducible mycelial morphology.^52^ The observations of the current study were also similar to the study described by Schrinner et al., (2020). Significant reduction in pellet diameter along with improved production of rebeccamycin titers synthesized by *L. aerocolonigenes* was observed.^53^

Customized morphology of the filamentous microbes could be achieved through the addition of inorganic microparticles to the culture.^54–57^ Various researchers used these strategies to achieve the suitable morphology for the improved production of desired compounds like polyketides, alcohols, different enzymes etc. from various filamentous fungi.^58–63^ Similar observations were reported by Driouch et al., 2011 where titanium silicate oxide was used as microparticle to alter the morphology of *A. niger.*^64^ Kuhl et al., 2021, observed 40 % reduction in pellet diameter along with boosted production of secondary metabolites by *S. lividans.*^65^ Holtman et al., 2017 studied the effect of SiO_2_ on the production of actinorhodin and achieved 85% increment in antibiotic production.^66^

Researchers also used medium engineering approach to obtain better yield of lipstatin, for instance, 1.98 mg/ml lipstatin production was reported by Kumar et al., 2011, whereas 3.92 mg/ml and 4.20 mg/ml production of lipstatin was observed in another study.^16, 67, 68^ However, no research has been published to date to improve the production of lipstatin or any other secondary metabolites produced by members of genus *Streptomyces* with the supplementation of natural precursors. Bule and Singhal (2009) reported the improved production of CoQ_10_ by the supplementation of carrot and tomato juice as natural precursors for *Pseudomonas diminuta* NCIM 2865.^69^

## Conclusion

In conclusion, present study demonstrated that the production of lipstatin is affected by pellet morphology. Small sized pellet with dispersed morphology provides higher yield of lipstatin. Along with pellet morphology, medium components also play important role in lipstatin production.

## Acknowledgements

The authors sincerely acknowledge the Central University of Haryana, Mahendergarh for necessary facilities. We also wish to acknowledge University Grant Commission (UGC), New Delhi for doctoral fellowship (597/(OBC) (CSIR-UGC NET DEC. 2016).

## Authors contribution

**Khushboo**: Conceptualization, Methodology, Formal analysis, Investigation, Writing – original draft. **Namrata Dhaka**: Writing – review & editing. **Kashyap Kumar Dubey**: Conceptualization, Methodology, Writing – review & editing, Supervision.

## Ethical Statements

The authors declare no conflict of interest.

Neither ethical approval nor informed consent was required for this study.

## Graphical abstract

**Fig. 1.**
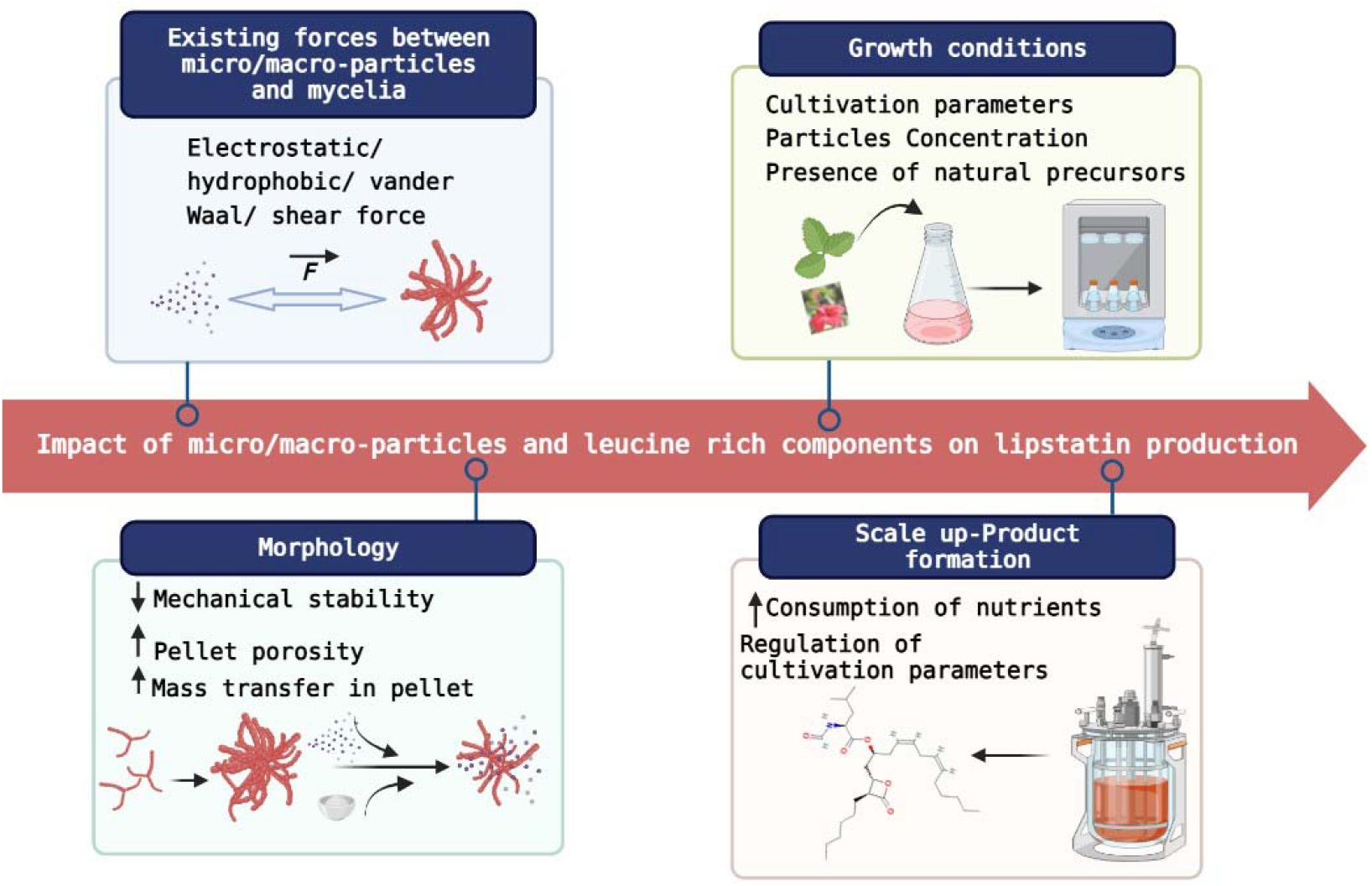

